# jackalope: a swift, versatile phylogenomic and high-throughput sequencing simulator

**DOI:** 10.1101/650747

**Authors:** Lucas A. Nell

## Abstract

High-throughput sequencing (HTS) is central to the study of population genomics and has an increasingly important role in constructing phylogenies. Choices in research design for sequencing projects can include a wide range of factors, such as sequencing platform, depth of coverage, and bioinformatic tools. Simulating HTS data better informs these decisions. However, current standalone HTS simulators cannot generate genomic variants under even somewhat complex evolutionary scenarios, which greatly reduces their usefulness for fields such as population genomics and phylogenomics. Here I present the R package `jackalope` that simply and efficiently simulates (i) variants from reference genomes and (ii) reads from both Illumina and Pacific Biosciences (PacBio) platforms. Genomic variants can be simulated using phylogenies, gene trees, coalescent-simulation output, population-genomic summary statistics, and Variant Call Format (VCF) files. `jackalope` can simulate single, paired-end, or mate-pair Illumina reads, as well as reads from Pacific Biosciences. These simulations include sequencing errors, mapping qualities, multiplexing, and optical/PCR duplicates. It can read reference genomes from FASTA files and can simulate new ones, and all outputs can be written to standard file formats. `jackalope` is available for Mac, Windows, and Linux systems.

## Introduction

High-throughput sequencing (HTS) is a cost-effective approach to generate vast amounts of genomic data and has revolutionized the study of genomes (Metzker, 2009). The combination of massive datasets, sequencing errors, and potentially complex evolutionary histories make bioinformatic pipelines an important aspect of research using HTS. Many bioinformatic tools exist, and new programs that are more accurate and computationally efficient are constantly being developed. Simulation of HTS data is needed to test these tools. Although there are many sequence simulators currently available (reviewed in Escalona, Rocha, & Posada, 2016), most have only rudimentary ways to generate variants from a reference genome. For example, both `Mason` (Holtgrewe, 2010) and `GemSIM` (McElroy, Luciani, & Thomas, 2012) place mutations randomly throughout genome sequences, the former based on per-site mutation rates and the latter on a set number of mutations per variant haplotype. In either case, simulations alone are not useful for researchers interested in simulating HTS data to test more complex evolutionary questions, such as (a) how demographic changes and selection interact to affect genome-wide diversity or (b) how incongruence between gene trees might affect reconstructed phylogenies.

Recently, pipelines have been developed that use multiple programs to simulate complex evolutionary histories and HTS on the resulting populations or species. `TreeToReads` (McTavish et al., 2017) simulates sequences along a single phylogenetic tree using `Seq-Gen` (Rambaut & Grassly, 1997) and generates Illumina reads using `ART` (Huang, Li, Myers, & Marth, 2011). The newer pipeline `NGSphy` (Escalona, Rocha, & Posada, 2018) uses `INDELible` (Fletcher & Yang, 2009) for more mutation options than `Seq-Gen` and can simulate along multiple gene trees via `SimPhy` (Mallo, Oliveira Martins, & Posada, 2015). These pipelines are quite powerful. However, they both require the installation of multiple programs to use them, and the vast array of features creates a steep learning curve when getting started. Additionally, the lack of integration between inner programs means that the entire process is not as computationally efficient as a single standalone program.

In the present paper I introduce `jackalope`, an R (R Core Team, 2019) package that combines the functionality of an HTS simulator with that of a phylogenomics simulator. Although `jackalope` can process output from coalescent simulators, it does not do coalescent simulations itself and assumes input files to be true. I chose the R platform because of its simple integration with C++ code via the `Rcpp` package (Eddelbuettel & François, 2011).

The interface with R allows the user greater flexibility, while the underlying C++ code improves memory use and speed. `jackalope` can read genomes from FASTA files or simulate them *in silico*. It can also create variants from the reference genome based on basic population-genomic summary statistics, phylogenies, gene trees, Variant Call Format (VCF) files, or matrices of segregating sites. These variants can be simulated based on any of several popular molecular-evolution models. `jackalope` simulates single, paired-ended, or mate-pair reads on the Illumina platform, as well as Pacific Biosciences (PacBio) reads. All information generated by `jackalope` can be output to standard file formats.

After outlining the methods, I compare the performance of `jackalope` to that of other popular programs. Lastly, I demonstrate the usefulness of `jackalope` through four usage examples.

## Features and methods

Most code is written in C++ and interfaces with R using the `Rcpp` package (Eddelbuettel & François, 2011). I used OpenMP to allow for parallel processing and the `PCG` family of thread-safe, pseudo-random number generators (O’Neill, 2014). I use alias sampling (Kronmal & Peterson, 1979; Walker, 1974) for all weighted sampling. Package `RcppProgress` provides the thread-safe progress bar (Forner, 2018). All input and output files can have `gzip` or `bgzip` compression, using the `zlib` and `htslib` (Li et al., 2009) libraries. Access to these libraries uses the R packages `zlibbioc` (Morgan, 2019) and `Rhtslib` (Hayden & Morgan, 2019) to improve portability.

There are two classes in `jackalope`, one that stores information on reference genomes (`ref_genome`), the other that stores information on sets of variants (`variants`). Both classes are `R6` (Chang, 2019) classes that wrap pointers to underlying C++ objects. Those C++ objects store all the information. Reference genomes are simply vectors of chromosome names and sequence strings. Variants are stored as nested vectors of mutation information. This can greatly reduce memory usage (unless the mutation density is high), and it provides the ability to output VCF files based on variants. I chose `R6` classes because they allowed me to make the pointers to the C++ objects private, thereby preventing users from accidentally manipulating them. Methods in `jackalope`’s classes allow the user to view and manipulate aspects of the C++ objects. Structuring the classes this way provides a high degree of flexibility and minimizes the chances of copying large objects in memory. An overview of how these classes are created and used to produce output files is shown in Figure 1.

**Figure 1:**
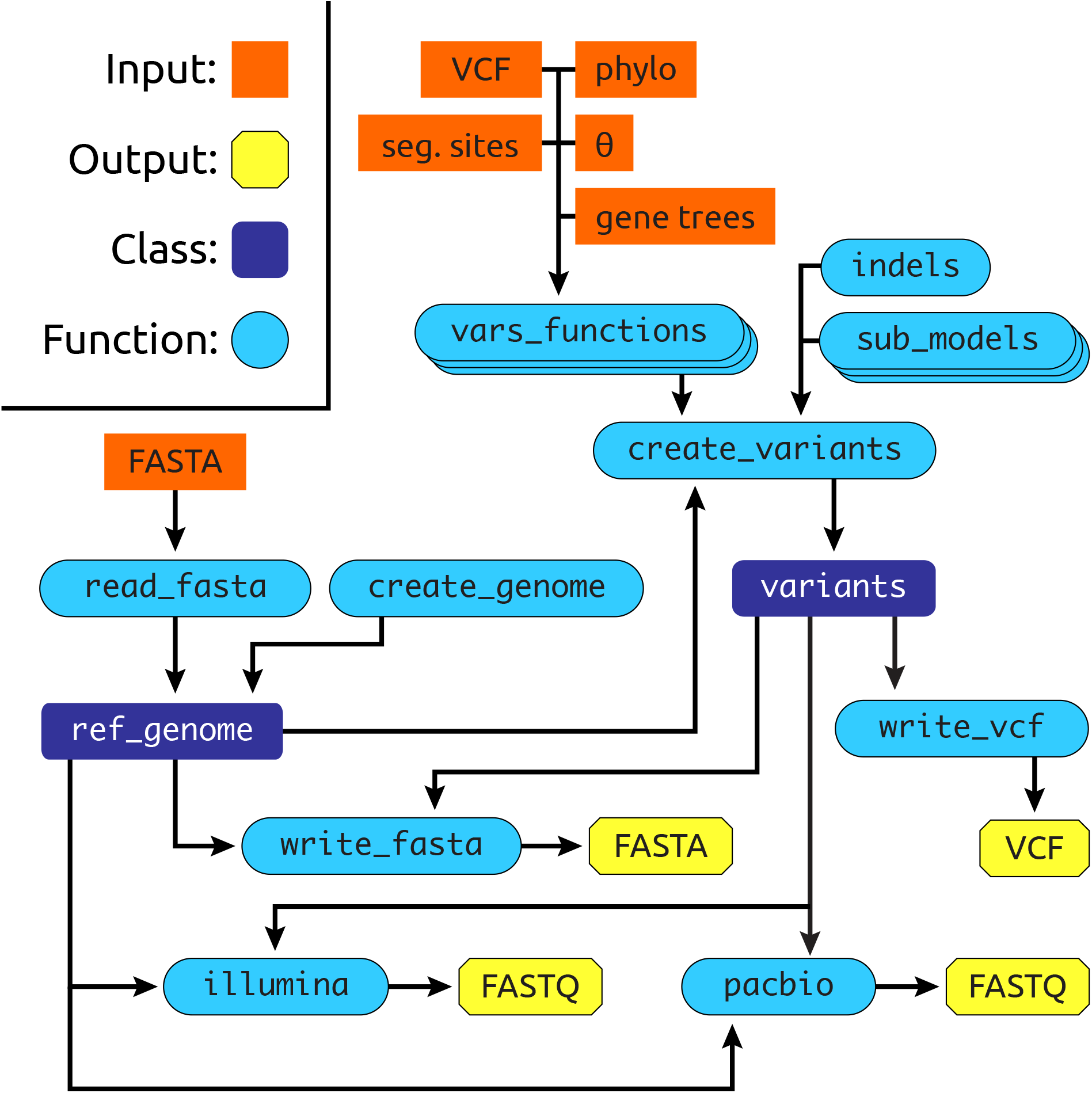
Overview of primary `jackalope` functions, classes, inputs, and outputs. Circles `sub_models` and `vars_functions refer` to multiple functions; see text for details. *θ* indicates the population-scaled mutation rate.

### Reference genomes

Haploid reference genomes can be generated from FASTA files, optionally with index files—created using `samtools faidx`—that speed up processing. If a reference genome is not available, one can be simulated based on given equilibrium nucleotide distributions. Chromosome lengths are drawn from a Gamma distribution with a mean and standard deviation provided by the user. The Gamma distribution was chosen because of its flexibility and due to evidence that it works well for chromosome sizes in diploid eukaryotes (Li et al., 2011).

### Creating variants

#### Methods

There are five ways to generate haploid variants from the reference genome, and information on all the methods can be found in the documentation (under `vars_functions`). The first two variant methods do not require phylogenetic information. First, a Variant Call Format (VCF) file can directly specify mutations for each variant. This method works using the `htslib` C library and is the only method that does not require any molecular-evolution information. Second, segregating sites from coalescent output can inform the locations of mutations and which variants have them. The user provides molecular evolution information, which is then used to sample for mutation types at each segregating site. Sampling is weighted based on each mutation type’s rate. The segregating-site information can take the form of (1) a coalescent-simulator object from the `scrm` (Staab, Zhu, Metzler, & Lunter, 2015) or `coala` (Staab & Metzler, 2016) package, or (2) a file containing output from a coalescent simulator in the format of the `ms` (Hudson, 2002) or `msms` (Ewing & Hermisson, 2010) program.

The remaining three variant methods use phylogenetic information. In the third method, phylogenetic tree(s) can be directly input from either `phylo` object(s) or NEWICK file(s). These can be a single species tree or one gene tree per chromosome. Fourth, users can pass an estimate for *θ*, the population-scaled mutation rate. A random coalescent tree is first generated using the `rcoal` function in the `ape` package (Paradis & Schliep, 2018). Then, its total tree length is scaled to be 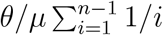 for *n* variants and an equilibrium mutation rate of *μ*. The last method allows for recombination by simulating along gene trees that can differ both within and among reference chromosomes. Similarly to simulations using coalescent segregating sites, gene trees can be from `scrm` or `coala` objects, or from `ms`-style output files.

In these last three “phylogenomic” methods, chromosomal sequences are evolved along phylogenetic trees (either species or gene trees). All chromosomes evolve independently. If recombination is included, multiple gene trees are used per chromosome, and chromosomal regions referred to by different gene trees evolve independently. Evolving independent chromosomal regions along phylogenetic trees starts by using the reference genome sequences as those for the root of the tree. Then, for each region and branch, the number of newly generated mutations is proportional to the branch length (see below for more details). These new mutations are then assigned to the daughter node on the tree. This process is repeated down the tree, from root to tips. In `jackalope`, mutation information is passed through the tree in such a way that no intermediate objects are created.

#### Molecular evolution

Both substitutions and indels can be simulated by `jackalope`. It is assumed that the substitution- and indel-producing processes are independent, so they are simulated separately to improve computational efficiency (Fletcher & Yang, 2009).

#### Substitutions

For substitutions, the following models can be employed: TN93 (Tamura & Nei, 1993), JC69 (Jukes & Cantor, 1969), K80 (Kimura, 1980), F81 (Felsenstein, 1981), HKY85 (Hasegawa, Kishino, & Yano, 1985; Hasegawa, Yano, & Kishino, 1984), F84 (Thorne, Kishino, & Felsenstein, 1992), GTR (Tavaré, 1986), and UNREST (Yang, 1994). If using the UNREST model, equilibrium nucleotide frequencies (*π*) are calculated by solving for *π***Q** = 0, where **Q** is the substitution rate matrix. This is done by finding the left eigenvector of **Q** that corresponds to the eigenvalue closest to zero. Each substitution model uses its own function, and the provided documentation (under `sub_models`) includes information to help users choose among them.

Substitution rates can also vary among sites. A random-sites, discrete-Gamma model is used, where sites’ rates are assumed to be independent of one another and are derived from a Gamma distribution split into *K* rate categories of equal probability (Yang, 2006). The Gamma distribution is constrained to have a mean of 1. A site’s overall rate is its nucleotide’s mutation rate multiplied by the “Gamma distance” associated with its rate category. The user can also specify a proportion of invariant sites.

For the phylogenetic methods, substitutions are simulated for each branch by first calculating the transition-probability matrix (*P*(*t*)) for each Gamma category based on the branch length. Substitutions are then sampled for each site using the *P*(*t*) matrix that coincides with the Gamma category at that site. This procedure has been used in multiple programs to capture the substitution process, including `Seq-Gen` (Rambaut & Grassly, 1997) and `INDELible` (Fletcher & Yang, 2009).

#### Indels

Information for insertions and deletions is provided separately, but the same information is needed for both. The first requirement is an overall rate parameter, which is for the sum among all nucleotides; indel rates do not differ among nucleotides. The relative rates of indels of different sizes can be provided in 3 different ways: First, rates can be proportional to exp(−*u*) for indel length *u* from 1 to the maximum possible length, *L* (Albers et al., 2010). Second, rates can be generated from a Lavalette distribution, where the rate for length *u* is proportional to [*uL*/(*L* − *u* + 1)]^−*a*^ (Fletcher & Yang, 2009). Third, relative rates can be specified directly by providing a length-*L* numeric vector of positive values. Indels up to 1 Mb are allowed.

Indels are generated along phylogenetic-tree branches using the *τ* -leaping approximation (Cao, Gillespie, & Petzold, 2006; Wieder, Fink, & Wegner, 2011) to the Doob–Gillespie algorithm (Doob, 1942; Gillespie, 1976). I chose the Doob–Gillespie algorithm because it only requires instantaneous rates rather than the calculation of a transition-probability matrix, making it easily used for indels (Yang, 2006). This method has been used in `INDELible` (Fletcher & Yang, 2009) and `Dawg` (Cartwright, 2005) to simulate indels. The Doob–Gillespie algorithm works by generating waiting times for each branch length that represent the time until the next mutation occurs somewhere on the sequence. Waiting times are generated from an exponential distribution with a rate equal to the sum of indel rates for all nucleotides in the sequence. Waiting times (and resulting mutations) are generated until the sum of all times is greater than the branch length.

The *τ*-leaping approximation improves efficiency by breaking the branch length into *τ*-sized chunks and using a Poisson distribution to generate how many mutations of each type occur in each chunk. The maximum value of *τ* is

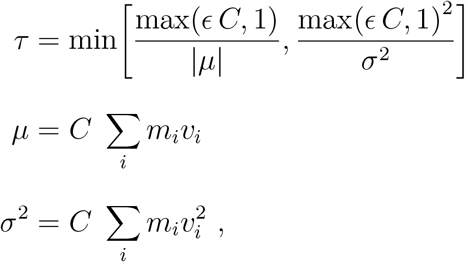

for an error control parameter *ϵ* (where 0 < *ϵ* ≪ 1), chromosome length *C*, mutation rate for one indel type *m_i_*, and effect on chromosome length for one indel type *v_i_* (Cao et al., 2006; Wieder et al., 2011). Increasing the error control parameter results in faster simulations that approximate the exact Doob–Gillespie algorithm less precisely. It can be changed by the user, but has a default value of 0.03 as recommended by Cao et al. (2006). The rate parameter for the Poisson distribution that generates the number of mutations over the entire chromosome across *τ* time units is *τCm_i_* for indel type *i*.

#### Simulating sequencing data

Both reference genomes and variants can be simulated for Illumina and PacBio sequencing. Chromosomes from which reads are derived are sampled with weights proportional to their length. If variants are provided, individual variants are sampled with equal probabilities by default. Alternatively, the user can specify sampling weights for each variant to simulate the library containing differing amounts of DNA from each. Both methods also allow for a probability of read duplication, which might occur due to PCR in either method and from optical duplicates in Illumina sequencing. `jackalope` can create reads using multiple threads by having a read “pool” for each thread and having pools write to file only when they are full. This reduces conflicts that occur when multiple threads attempt to write to disk at the same time. The size of a “full” pool can be adjusted, and larger sizes should increase both speed and memory usage. Illumina reads can be single, paired-ended, or mate-pair, and I used methods based on the `ART` program (Huang et al., 2011). PacBio read simulation is based on `SimLoRD` (Stöcker, Köster, & Rahmann, 2016). Each was re-coded in C++ to more seemlessly integrate into `jackalope`. Function inputs emulate the program they were based on, and users are encouraged in each function’s documentation to also cite `ART` or `SimLoRD`.

#### Writing output

Reference genomes and variants can be written to FASTA files; the latter has a separate file for each variant. Variant information can also be written to VCF files, where each variant can optionally be considered one of multiple haplotypes for samples with ploidy levels > 1. Simulated gene trees can be written to `ms`-style output files, and phylogenies can be written to NEWICK files using `ape∷write.tree`. All sequencing reads are output to FASTQ files.

## Validation and performance

I validated `jackalope` by conducting a series of simulations for variant creation and Illumina and PacBio sequencing. Using known inputs, I compared predicted to observed values of outputs, and `jackalope` conformed closely to expectations (Supporting Information Figures S1–S13).

I compared the performance of `jackalope` to existing programs on a MacBook Pro running macOS Catalina (version 10.15) with a 2.6GHz Intel Core i5 processor and 8 GB RAM. Elapsed time and maximum memory used were measured by using the GNU `time` program (`/usr/bin/time` from the terminal). For each test, 5 runs of each program were tested in random order. In the text below, numbers presented are means from these 5 tests unless otherwise stated.

### Creating variants

For creating variants, I compared `jackalope` to `INDELible` (version 1.03; Fletcher & Yang, 2009). I chose `INDELible` because it is used by `NGSphy` and because it can generate sequences with both substitutions and indels based on phylogenetic trees. I tested each program by having them simulate a 2 Mb and 20 Mb genome split evenly among 20 chromosomes, then generate 8 variants from that genome using phylogenetic trees produced by the command line version of `scrm`. I ran separate simulations with the trees scaled to maximum tree depths of 0.1, 0.01, and 0.001. I used the HKY85 substitution model with a transition rate of 2.5 substitutions per site per unit of time (where one unit of time is equal to a branch length of 1), and a transversion rate of 1. Relative frequencies of nucleotides were 0.4, 0.3, 0.2, and 0.1 for T, C, A, and G, respectively. For among-site variability in substitution rates, I used (1) a discrete Gamma distribution with 10 categories and a shape parameter of 0.5, and (2) an invariant-site rate of 0.25. The total rate for insertions was 0.1 per site per unit time, with relative rates derived from a Lavalette distribution with *L* = 541 and *a* = 1.7. Deletions had the same total and relative rates. Output was written to FASTA files. `jackalope` was tested for 1 and 4 threads, but `INDELible` for only 1 because it cannot use multiple threads.

`jackalope` consistently outperformed `INDELible` in terms of speed, although this difference was less drastic when there were more mutations: for the larger genome size and greater maximum tree depth (Figure 2). This is somewhat offset by the fact that using more threads has a greater effect at larger genome sizes. Memory usage was greater in `jackalope` than `INDELible`. With low mutation numbers, this difference was due to the overhead associated with loading R. For a 2 Mb genome and 0.001 max tree depth, `jackalope` used 94.3 MB memory, while `INDELible` used 20.7 MB. An R script that simply printed an empty string used ~ 55 MB memory, and another that only loaded `jackalope` used ~ 70 MB. With greater numbers of mutations, the difference was due to `jackalope` using 64-bit integers to store positions instead of 32-bit. In the most extreme example (20 Mb genome and 0.1 max tree depth), `jackalope` used 768.3 MB memory and `INDELible` used 429.3 MB. I chose to use 64-bit integers in `jackalope` to allow chromosomes to exceed 2 Gb in length. Memory usage increased by only 31.7kB when `jackalope` used 4 threads.

**Figure 2:**
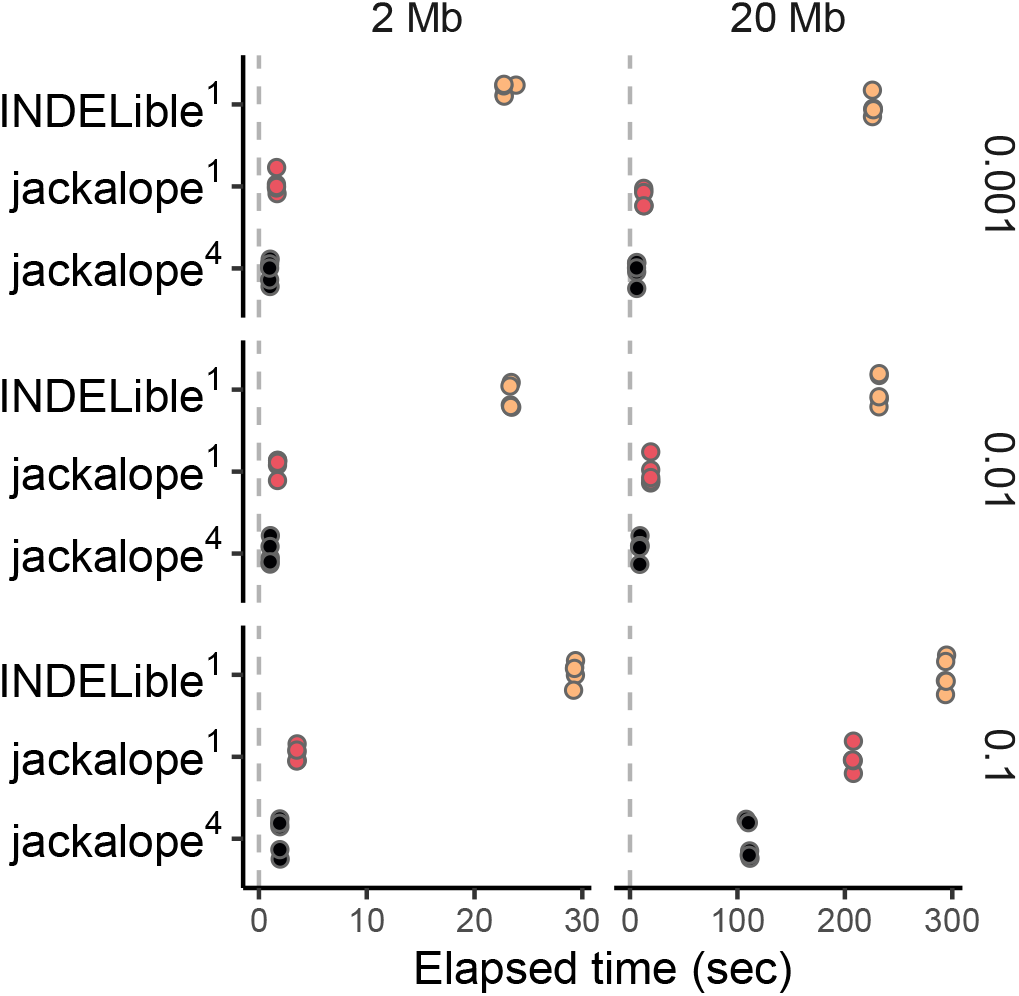
Performance comparison between `jackalope` and `INDELible` in generating variants from a reference genome. Sub-panel columns indicate the size of the genome from which variants were generated. Sub-panel rows indicate the maximum tree depth of the phylogeny used for the simulations. Superscripts in the y-axis labels indicate the number of threads used.

### Generating sequencing reads

For both Illumina and PacBio read generation in `jackalope`, I compared their performance to the programs they were based on: `ART` (version 2.5.8; Huang et al., 2011) and `SimLoRD` (version 1.0.3; Stöcker et al., 2016), respectively. `ART` has the added benefit of being used for both `NSGphy` and `TreesToReads`. All these tests consisted of reading a 2 Mb genome from a FASTA file, simulating reads, and writing them to FASTQ files. No ALN or SAM/BAM files were written by any of the programs. For Illumina sequencing, I generated 2 × 150 reads from the HiSeq 2500 platform, and tests were conducted for 100 thousand, 1 million, and 10 million total reads. For PacBio sequencing, I generated reads using default parameters, and tests were conducted for 1, 10, and 100 thousand total reads. `jackalope` was tested for 1 and 4 threads, but neither `ART` nor `SimLoRD` allowed the use of more than 1.

Again, `jackalope` outperformed both `ART` and `SimLoRD` in terms of speed (Figure 3A–B). Multithreading was most useful when many reads were generated: For at least 10M Illumina reads and at least 100k PacBio reads, using 4 threads reduced the elapsed time for jobs by ~ 50%. All three programs used very little memory: Average memory usage was 9.3 MB for `ART` and 60.9 for `SimLoRD`. `jackalope` averaged 73.7 MB for Illumina reads and 102.0 for PacBio reads. Memory usage increased only slightly with increasing read numbers in `jackalope:` It increased 0.6 MB across the range of Illumina reads and 3 MB across the PacBio range. Over the Illumina range, memory usage by `ART` slightly decreased (−0.05 MB). `SimLoRD` memory usage increased over its range (8 MB). Using 4 threads in `jackalope` had little effect on memory usage (3.2 MB extra for Illumina reads, 6.3 MB extra for PacBio).

**Figure 3:**
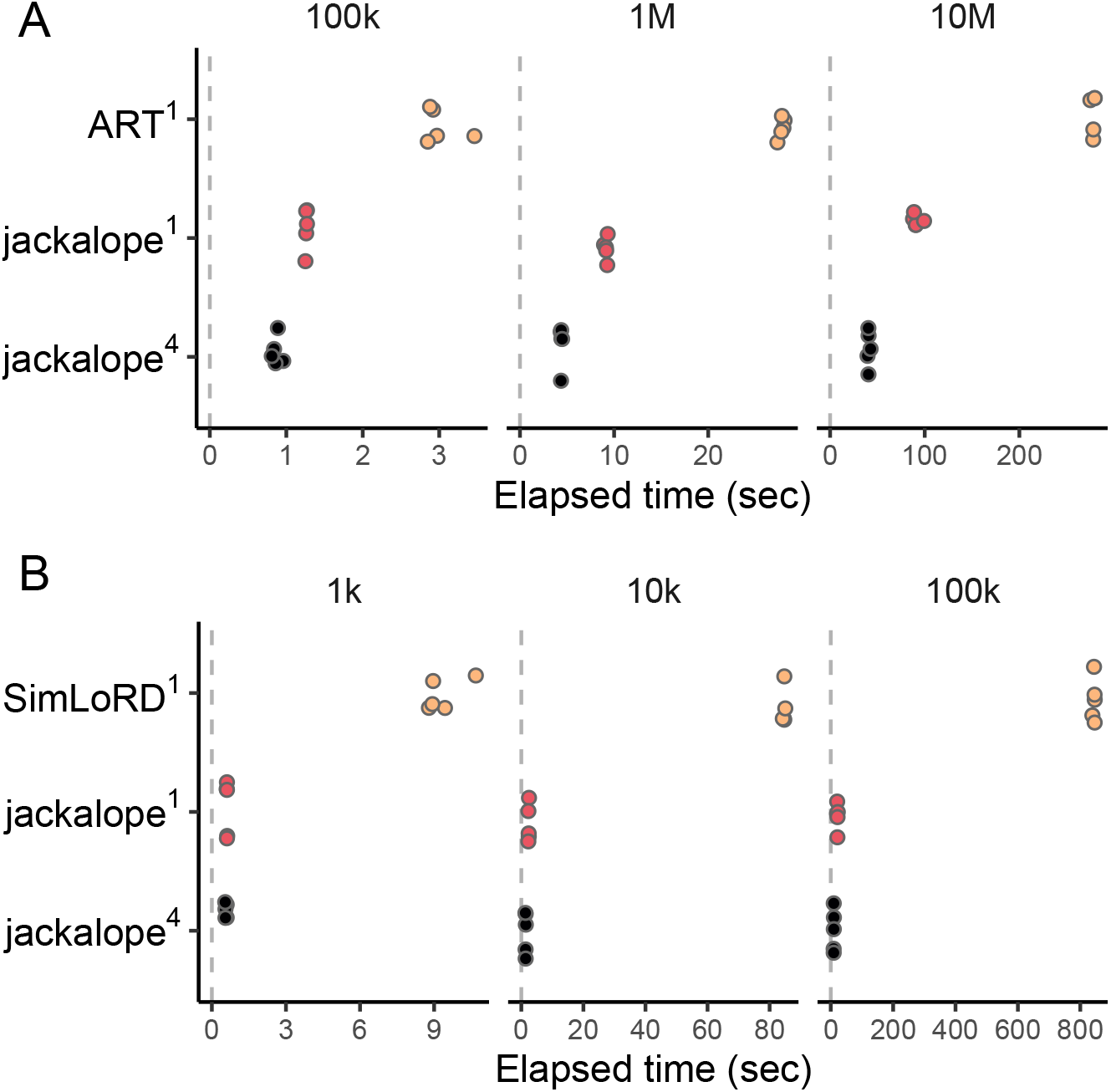
Performance comparison between `jackalope` and (A) `ART` in generating Illumina reads and (B) `SimLoRD` in generating PacBio reads. Sub-panel columns indicate the number of reads generated. Superscripts in the y-axis labels indicate the number of threads used.

### Pipeline from reference to reads

I lastly compared the performance of `jackalope` to that of `NGSphy` (version 1.0.13; Escalona et al., 2018) for the following pipeline: generating a reference genome, simulating variants from gene trees, and producing Illumina reads from the variants. For the first two steps, I used the same parameters as I did for the comparison to `INDELible`. I also used the same parameters for the last step, except that the haploid variants were used for sequencing reads instead of a reference genome. I generated 8 variants with 20 chromosomes by creating a phylogenetic tree with 160 tips and assigning every 20 tips to the same individual using the naming scheme necessary for `NGSphy`. I divided the larger tree by chromosome and passed these separate trees to `jackalope` by writing them to a `ms`-style output file. Output was written to both FASTA and FASTQ files. Both `NGSphy` and `jackalope` were tested using both 1 and 4 threads.

`jackalope` was faster than `NGSphy` for all the performance tests (Figure 4A). The speed advantage of `jackalope` was greatest at low mutation densities and when the number of reads generated was higher. The effect of multithreading in `jackalope` increased with increasing read numbers, genome sizes, or maximum tree depth. These results were both consistent with those from the separate performance tests with `INDELible` and `ART`, the programs that make up the `NGSphy` pipeline. Attempts at multithreading in `NGSphy` had no effect on its performance: Calling for 4 threads resulted in 0.7 % more elapsed time and never caused total CPU time to exceed elapsed time. Thus multithreading in `NGSphy` seems to take place in another portion of the pipeline that is not apparent from its documentation. For this reason, I have grouped results from both the 1- and 4-thread calls to `NGSphy` together in Figure 4 and in the text below.

**Figure 4:**
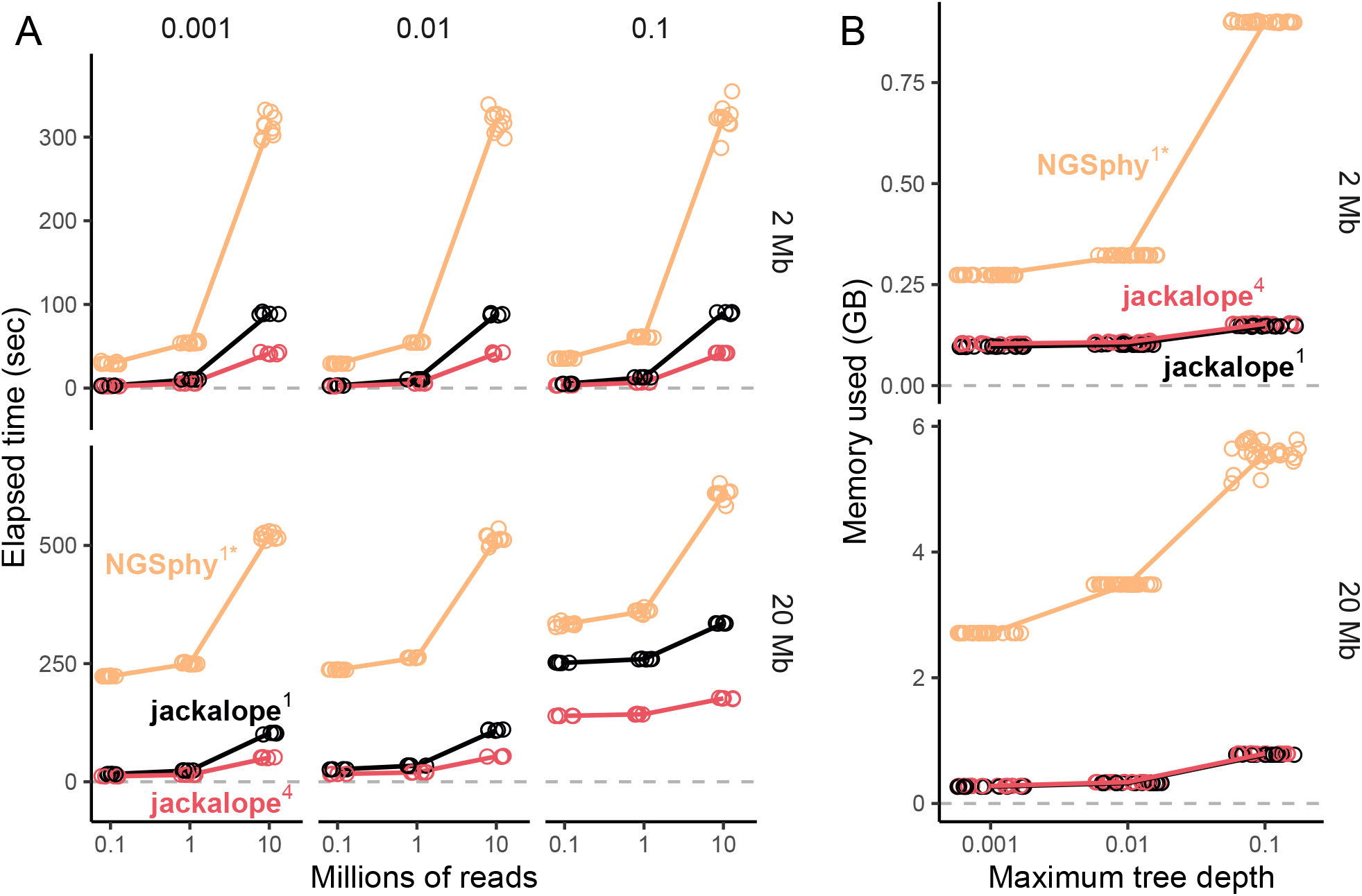
(A) Speed and (B) memory usage comparison between `jackalope` and `NGSphy` in generating variants from a reference genome, then producing Illumina reads from those variants. In (A), sub-panel columns indicate the maximum tree depth of the phylogeny used for the simulations. In both plots, sub-panel rows indicate the size of the genome from which variants were generated, and color indicates the program used; superscripts in the color labels indicate the number of threads used. The * highlights where results were combined for calls to `NGSphy` with 1 and 4 threads; see text for details.

Throughout the tests, `NGSphy` used significantly more memory than `jackalope` (Figure 4B). This was particularly true for the 20 Mb genome at maximum tree depths of 0.001 and 0.01, where `NGSphy` consumed 892 % and 947 % more memory, respectively. The number of reads had little effect on memory usage: When generating 10M versus 100k reads, `jackalope` consumed 0.02 % less memory, and `NGSphy` used 0.6 % less.

## Example usage

This section provides brief examples of how `jackalope` can be used to generate sequencing data that can inform some common sampling decisions for HTS studies. These examples only show how to produce the output from `jackalope`, as a review of different bioinformatic programs that would process these data is beyond the scope of the present manuscript.

For an example reference assembly, I used the *Drosophila melanogaster* assembly (version 6.27) downloaded from `flybase.org` (Thurmond et al., 2018). After downloading, I read the compressed FASTA file, filtered out scaffolds by using a size threshold, and manually removed the sex chromosomes using the following code:

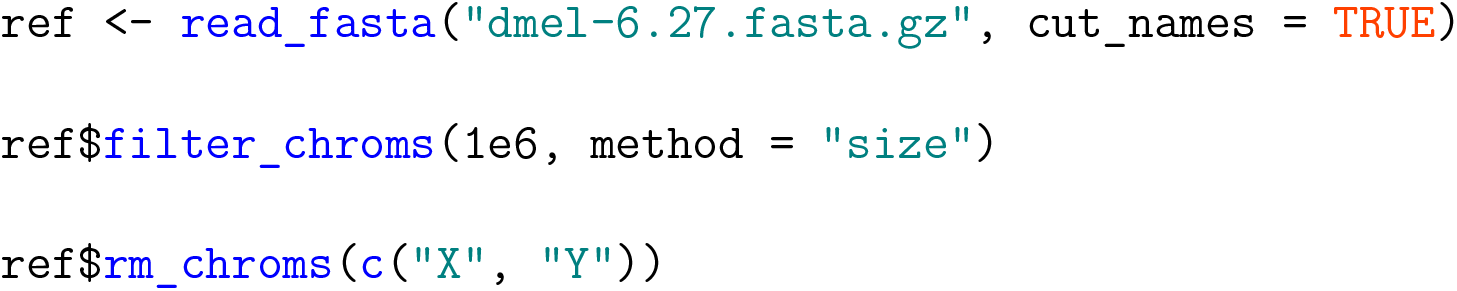

For mutation-type information, I used the JC69 substitution model, an insertion rate of `2e-5` for lengths from 1 to 10, and a deletion rate of `1e-5` for lengths from 1 to 40.

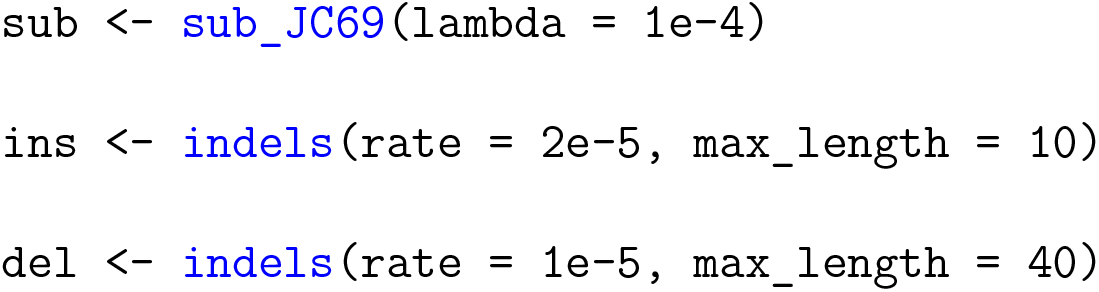

### Assembling a genome

The example here produces FASTQ files from the known *D. melanogaster* reference assembly that could test strategies for how to assemble a similar genome using HTS data. The strategies the example code below could test would be (1) a PacBio-only assembly, (2) a PacBio-only assembly followed by polishing using Illumina reads, and (3) an explicit hybrid assembly.

To generate data to test the first, PacBio-only strategy, I ran the following for two Single Molecule, Real-Time (SMRT) cells from the PacBio Sequel system (with the file `pacbio_R1.fq` as output):

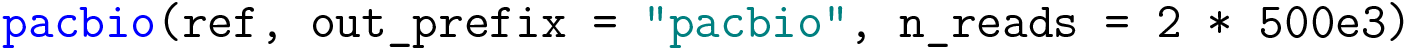

For either of the last two strategies, I used 1 SMRT cell of PacBio sequencing and 1 lane (~ 500 million reads) of 2 × 100bp Illumina sequencing on the HiSeq 2500 system (the default Illumina system in `jackalope`):

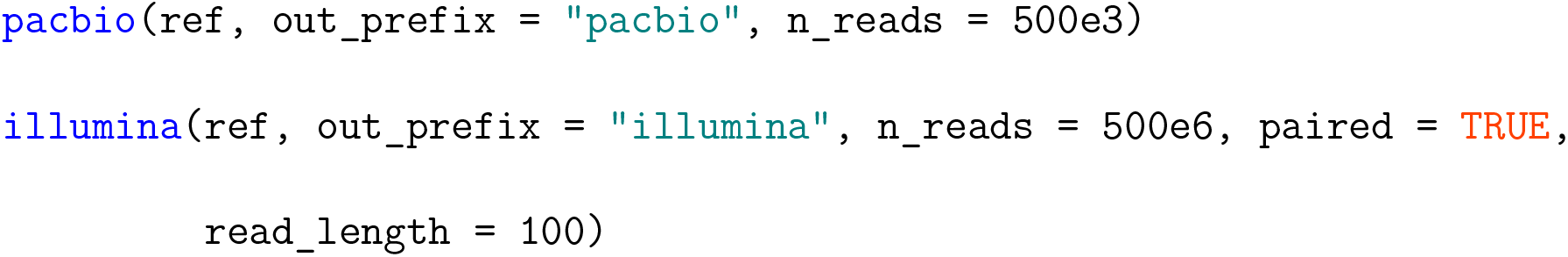

These data could then be used to compare genome assembly performance between the strategies above, or between programs within a given strategy. Extensions of these tests include adjusting sequencing depth, sequencing platform, and error rates.

### Estimating divergence between populations

Here, I will demonstrate how to generate pooled population-genomic data (Pool-seq) that could be used to estimate the divergence between two populations. Before starting `jackalope`, I used the `scrm` package (Staab et al., 2015) to conduct coalescent simulations that will generate segregating sites for 40 haploid variants from the reference genome (object `ssites`). Twenty of the variants are from one population, twenty from another. The symmetrical migration rate is 100 individuals per generation, and the population-scaled mutation rate (*θ* = 4*N*_0_*μ*) is 10. The `scrm` command for this is `40 5 -t 10 -I 2 20 20 100`.

Using the previously created objects for mutation-type information and the vars_ssites function that checks and organizes the coalescent output object, I created variants from the reference genome:

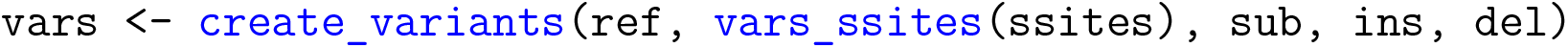

Lastly, I simulated 1 lane of 2 × 100bp Illumina HiSeq 2500 sequencing. In this case, each individual variant’s reads were output to separate FASTQ files. I also wrote the true variant information to a VCF file.

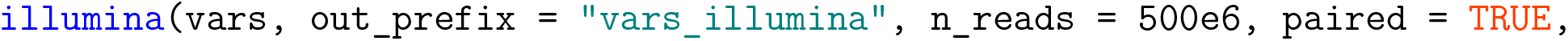

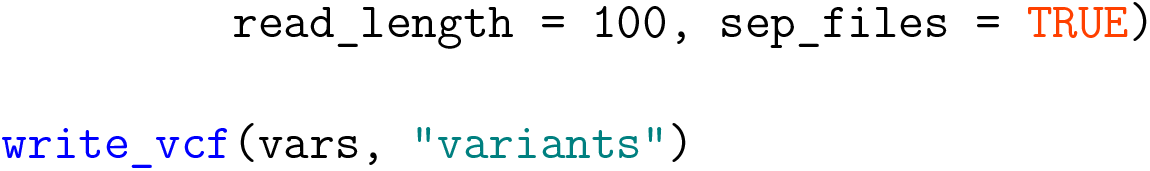

The *F_ST_* calculated from the resulting VCF file could then be compared to output from various programs to inform which works best in a particular case. For uncertain population parameters (e.g., migration rates), output from multiple calls to `scrm` varying the parameter of interest could be input to the `jackalope` pipeline above to identify the conditions under which one program might have an advantage over another.

### Constructing a phylogeny

#### From one phylogenetic tree

This section shows how `jackalope` can generate variants from a phylogeny, then simulate sequencing data from those variants to test phylogeny reconstruction methods. First, I simulated a coalescent phylogeny of 10 species (object `tree`) using `ape∷rcoal(10)` (Paradis & Schliep, 2018). Function `vars_phylo` organizes and checks the `tree` object, and including it with the mutation-type information allowed me to create variants based on this phylogeny:

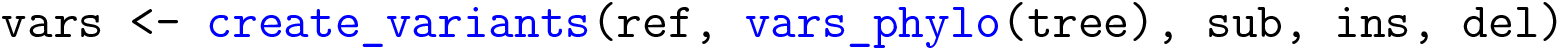

I then generated data for 1 flow cell of 1 × 250bp sequencing on an Illumina MiSeq v3, where `variant_barcodes` is a character string that specifies the barcodes for each variant. I also wrote the true phylogenetic tree to a NEWICK file.

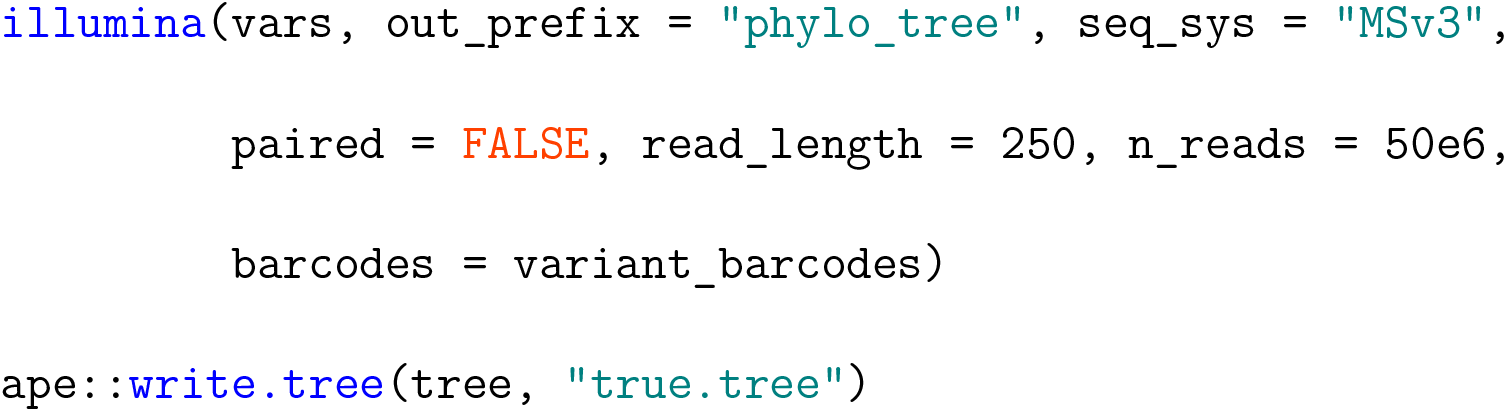

The true phylogenetic tree would then be compared to the final tree output from the program(s) the user chooses to test.

#### From gene trees

Similar to the section above, the ultimate goal here is to test phylogeny reconstruction methods. The difference in this section is that instead of using a single, straightforward phylogeny, I use multiple gene trees per chromosome. In the species used in these simulations, species 3 diverged from 1 and 2 at *t* = 1.0, where *t* indicates time into the past and is in units of 4*N*_0_ generations. Species 1 and 2 diverged at *t* = 0.5. I assume a recombination rate of 1/(4*N*_0_) recombination events per chromosome per generation. There are 4 diploid individuals sampled per species. I used `scrm` to simulate the gene trees (command `24 1 -T -r 1 1000 -I 3 8 8 8 0 -ej 1.0 3 1 -ej 0.5 2 1` for a chromosome size of 1000), producing the object `gtrees`. As for the other variant-creation methods, function vars_gtrees checks and organizes information from the `gtrees` object.

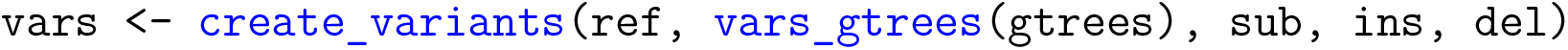

Next I generated data for 1 flow cell of 2 × 250bp sequencing on an Illumina MiSeq v3, with barcodes in the object `variant_barcodes`. I also wrote the true gene trees to a `ms`-style file.

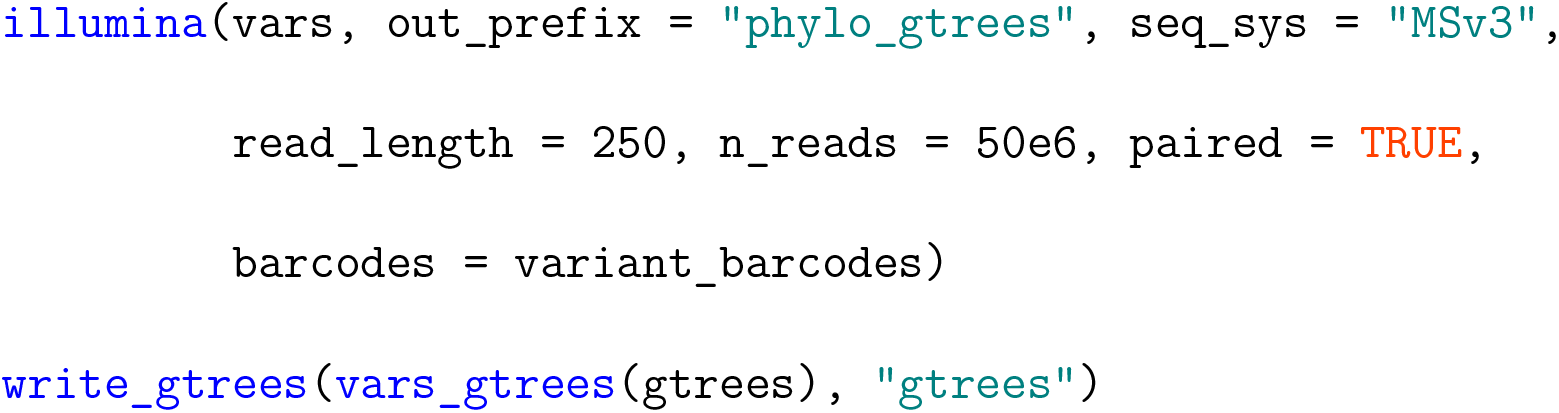

Topologies of the gene trees would then be compared to final phylogenies output from software the user is interested in testing. Varying recombination rates or adding gene flow after separation of species would be natural extensions of these simulations.

## Conclusion

`jackalope` outperforms popular stand-alone programs for phylogenomic and HTS simulation and combines their functionalities into one cohesive package. Although it does not provide the in-built power of a full pipeline like `NGSphy`, simulations using `jackalope` are simpler to implement and more efficient computationally. As part of a larger pipeline that includes coalescent simulations or other methods that guide patterns of mutations, `jackalope` should inform research design for projects employing HTS, particularly those focusing on population genomics or phylogenomics. Output from `jackalope` will help develop more specific sequencing goals in funding applications and estimate the power of a given sequencing design. Furthermore, `jackalope` can be used to test bioinformatic pipelines under assumptions of much more complex evolutionary histories than most current HTS simulation platforms allow.

## Data Accessibility

`jackalope` is open source, under the MIT license. The stable version of `jackalope` is available on CRAN (https://CRAN.R-project.org/package=jackalope), and the development version is on GitHub (https://github.com/lucasnell/jackalope). The documentation can be found at https://jackalope.lucasnell.com. The version used in this manuscript was 1.0.0. Code for the example usage, validation, and performance is available on GitHub at https://github.com/lucasnell/jlp_ms.

## Author Contributions

L.A.N. conceived and designed the project, wrote the code, and wrote the manuscript.

