## Supporting Information for "jackalope: a swift, versatile phylogenomic and high-throughput sequencing simulator"

Lucas A. Nell[1]

[1]Department of Integrative Biology, University of Wisconsin–Madison

This document provides diagnostics on creating variants, Illumina sequencing, and Pacific Biosciences (PacBio) sequencing in `jackalope`. The overall conclusion is that the simulations work as expected.

### 1 Creating variants

This tests that when creating variants, `jackalope` produces substitutions, indels, and among-site variation that fit expectations. It lastly checks that when simulating along a phylogeny, the number of differences between individual variants is proportional to the phylogenetic distance between them.

To conduct these tests, `jackalope` is compiled with `__JACKALOPE_DIAGNOSTICS` defined inside the `src/jackalope_config.h` file. This causes `jackalope` to print information about mutations as they are produced and about among-site variation as the rates are generated. In most of the sections below, I captured this console output to describe the observed output from `jackalope`.

The reference genome used in this section is displayed below:

```
## < Set of 20 chromosomes >
## # Total size: 2,000,000 bp
##   name                          chromosome                          length
## chrom0     GGTAGTCATGTTAGGTTACGAGGTT...TCGTAGACATGTAGCCCACACTTTCA    100000
## chrom1     GCTCCGATCTGTCGCCCCATTGGTC...CGACCCATATGGTTCGGTAGTCGGCG    100000
## chrom2     TGTTAGAAGGTCCGACCGAAATTAG...CACCCGGAACTATTCTAGCCTGCAAG    100000
## chrom3     TAGTGTGAGATGGTCATGTTACGTA...CCGCTACGACCAGAAGGAAACCGTCC    100000
## chrom4     CGGTTTGCCAGGGGGACATCCATAG...CGGTAGAGCACCGTCGGTCCGTTTAC    100000
## ...        ...                                                           ...
## chrom16    TGACCTCGCACGAATGTTGGCATCC...GGGGACATGTTCGAGCGCACAAGTCT    100000
## chrom17    AGAGGGGTGCTTCGCCTTAGGATCG...CGTGGCTTCGGATTTTTGGCATGTAA    100000
## chrom18    TATCCCAGGTAAAACATCGATGTAA...GCCCTTGAGATTATCTGAGGCCTGAT    100000
```

```
## chrom19    TCTCAAGTATTATGACGTTCGGCTA...TTCATTTCAAGTTCAGTGAAGTCACC    100000
```

#### 1.1 Substitution rates

This simulates just substitutions along a phylogeny of 2 variants, with a branch length of 0.1 and with the following substitution rate matrix:

```
##           T      C      A      G
##    T -0.400  0.050  0.150  0.200
##    C  0.025 -0.375  0.150  0.200
##    A  0.050  0.100 -0.290  0.140
##    G  0.050  0.100  0.105 -0.255
```

I examined the substitutions from the output object, rather than from console diagnostics, because it was simpler this way since there was no among-site variation. I counted the number of each type of mutation (corresponding to all non-diagonal elements in the rate matrix) for each chromosome and variant. I compared these to the predicted substitution counts, which were the product of the chromosome size and the transition-probability matrix for the branch length of the phylogeny.

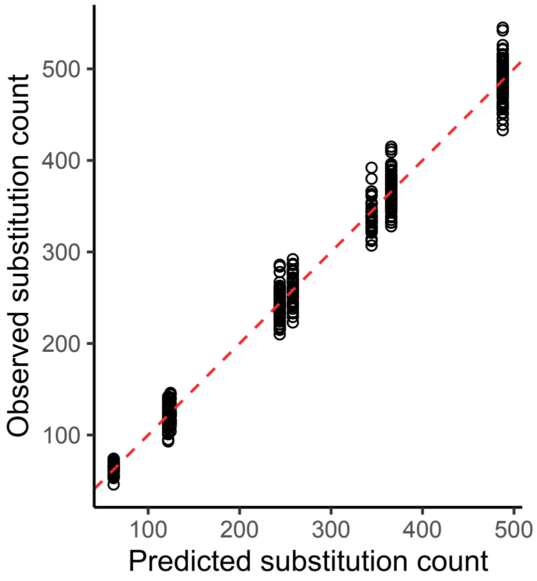

**Figure S1:** Observed versus predicted substitutions for simulation of 2 variants along a phylogeny.

#### 1.2 Indel rates

In this section, I first set the deletion rate to zero and the insertion rate to 0.2. In the second set of simulations, the insertion rate was zero and the deletion rate was 0.3.

I used the diagnostic console output from `jackalope` to examine how each round of indel production compared to expectations. For each "$\tau$-leap" (see Methods in main text for details), the console output the chromosome size, $\tau$, and the number of indels of each size that were produced. This was important to do for each round because the chromosome size changed each time. The predicted number of insertions of size $u$ was the product of the overall insertion rate, $\tau$, chromosome size, and relative rate for size $u$. The same goes for deletions.

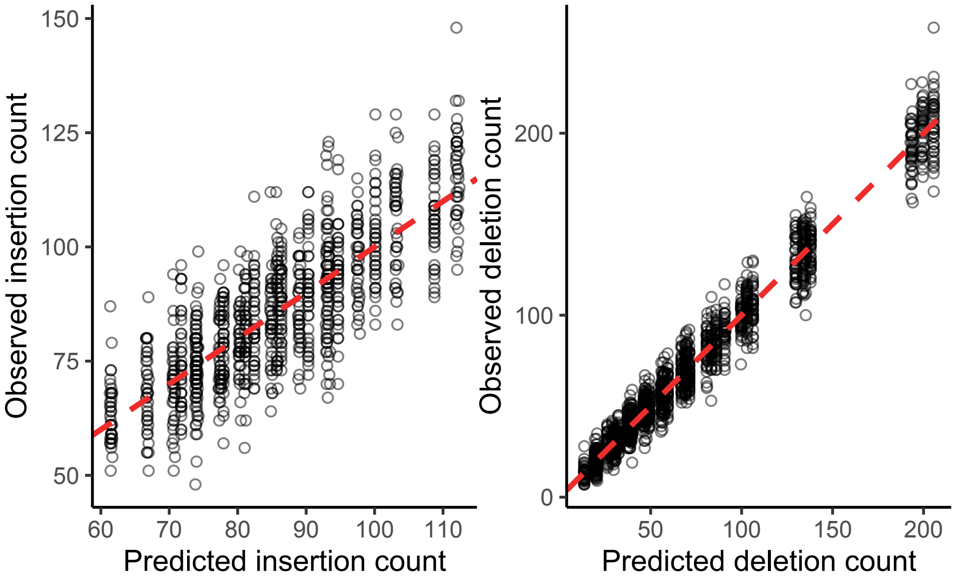

**Figure S2:** Observed versus predicted insertions (left) and deletions (right) for simulation of 2 variants along a phylogeny.

#### 1.3 Among-site variation

In this set of simulations, I included only substitutions because indels are not affected by among-site variation. I used the same substitution rate matrix as in the "Substitution rates" section. For the discrete Gamma distribution used to create among-site variation, I used a shape value of 0.5 and split the distribution into 5 categories.

I used the diagnostic console output for this section. Each time a substition was generated, the console output the position, Gamma-distribution category (0 to 4 in this case), the original nucleotide, and the new nucleotide. It also output when switching chromosomes, so I could match substitutions with chromosomes. I counted the number of mutations per chromosome, rate category, and type of substitution. I compared these to predicted counts, which were the product of the chromosome size, number of variants, and the transition-probability matrix for the branch length of the phylogeny.

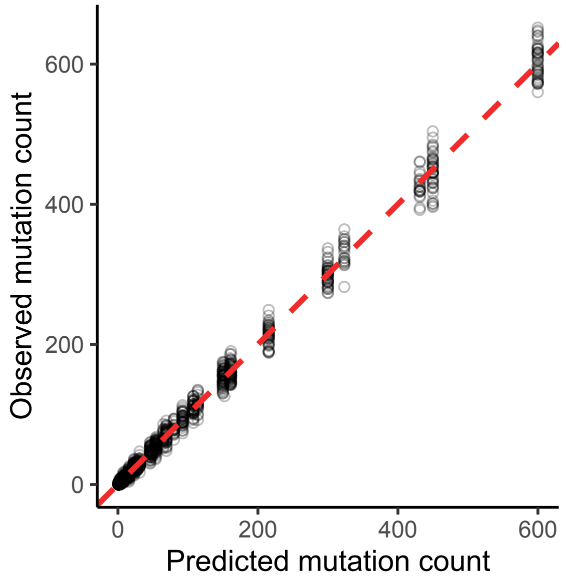

**Figure S3:** Observed versus predicted number of mutations, based on each position's mutation rate modifier from a discrete Gamma distribution. The dashed line is the 1-to-1 line.

I also checked that the number of sites were approximately equal for each category. This should be true because each rate category represents an equal area under the probability density curve for the Gamma distribution. The diagnostic output included when rates were created so that I could directly estimate how many sites belonged in each rate category.

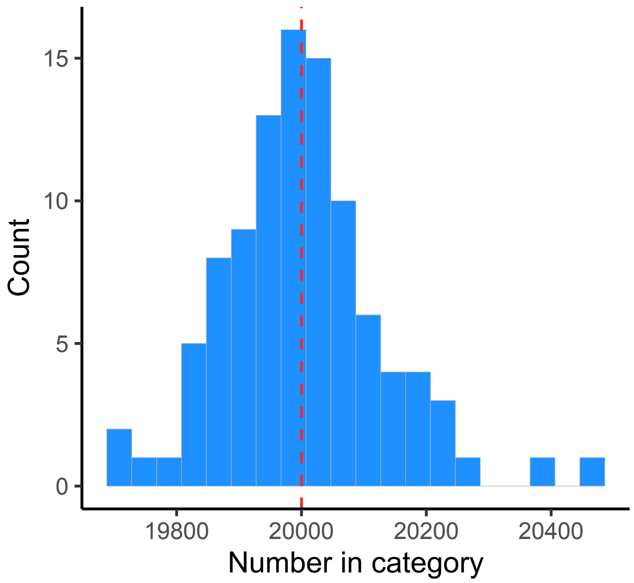

**Figure S4:** Observed number of sites in each rate category. The red line is the predicted value for all categories.

#### 1.4 Phylogeny

For this section, I simulated 10 variants along the following phylogenetic tree:

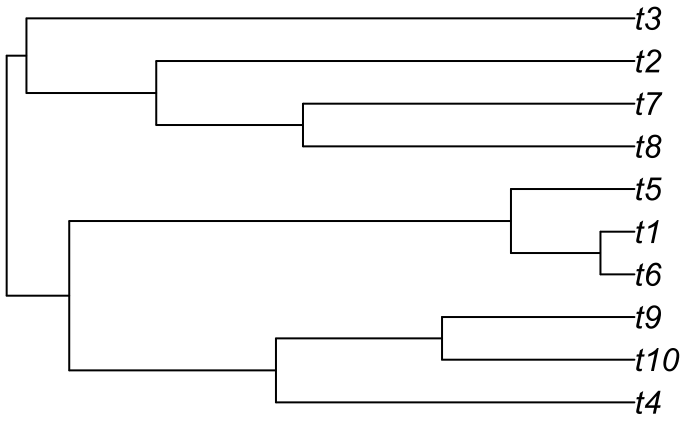

**Figure S5:** Phylogeny of 10 simulated variants.

I used only substitutions in the simulations, to make matching mutations between variants more straightforward.

I used the diagnostic console output for this section. Each time a substition was generated, the console output the position, Gamma-distribution category (always 1 in this case), the original nucleotide, and the new nucleotide. It also output when switching to a new edge on the phylogeny, so I could match substitutions with branch lengths. I counted the number of mutations per tree edge and compared these to predicted counts, which were the product of the chromosome size and the transition-probability matrix for the branch length.

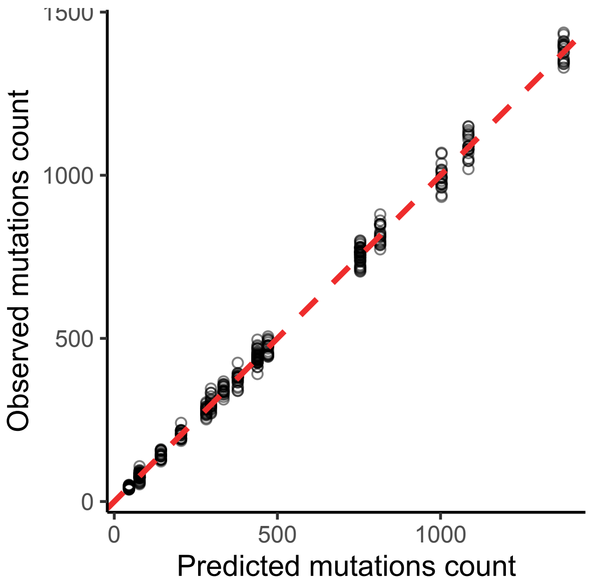

**Figure S6:** Observed versus predicted numbers of unshared mutations between 10 variants created by simulating along a phylogeny. Each point represents the number of unshared mutations between two variants for one of the 20 chromosomes. The dashed line is the 1-to-1 line.

### 2 Illumina

This section tests that the Illumina sequencer produces output that fits our expectations.

#### 2.1 Testing mismatches and quality scores:

Mismatches and quality scores are simulated the same as in ART. First, qualities profiles indicate the frequency of quality scores by position on read. Then, the probability of a mismatch for quality $q$ is $10^{-q/10}$. I am testing the accuracy of the output from `illumina` by starting with a reference genome consisting of 4 sequences, each containing a single nucleotide repeated 100,000 times:

```
## < Set of 4 chromosomes >
## # Total size: 400,000 bp
##   name                          chromosome                              length
```

```
## chrom0     TTTTTTTTTTTTTTTTTTTTTTTTT...TTTTTTTTTTTTTTTTTTTTTTTTTT    100000
## chrom1     CCCCCCCCCCCCCCCCCCCCCCCCC...CCCCCCCCCCCCCCCCCCCCCCCCCC    100000
## chrom2     AAAAAAAAAAAAAAAAAAAAAAAAA...AAAAAAAAAAAAAAAAAAAAAAAAAA    100000
## chrom3     GGGGGGGGGGGGGGGGGGGGGGGGG...GGGGGGGGGGGGGGGGGGGGGGGGGG    100000
```

Then I simulated 100,000 100-bp reads from the HiSeq 2500 platform ten times. Reads were matched with the sequence they derived from based on which nucleotide comprised the majority of the read. I counted mismatches as nucleotides that differ from the majority nucleotide. Qualities are directly measured from the output FASTQ. Both mismatches and qualities are aggregated by their position on the read, and compared to expectations as calculated from the profile file.

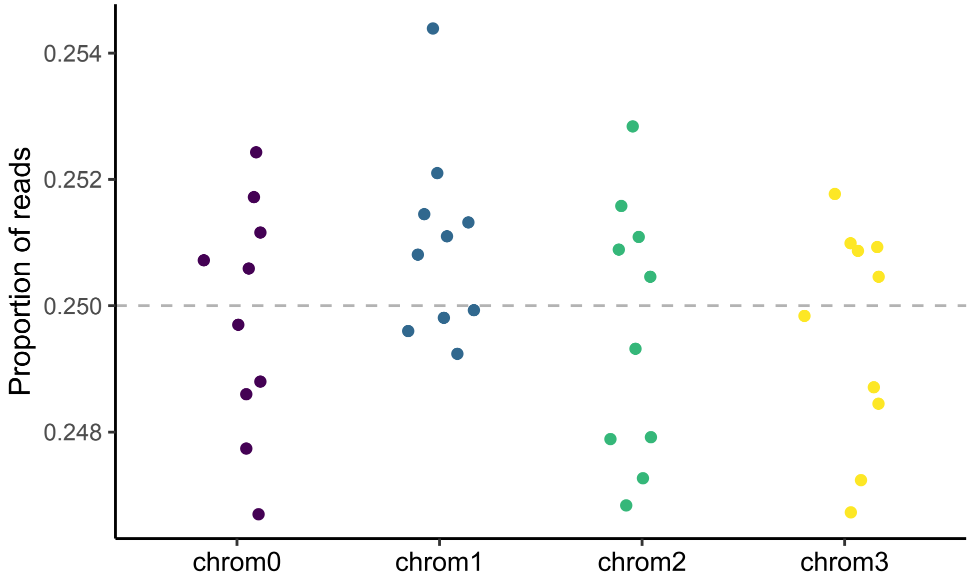

**Figure S7:** The proportion of reads derived from each reference genome sequence. Since all sequences were the same size, reads were expected to be generated from each sequence with equal probability (indicated by the horizontal gray dashed line).

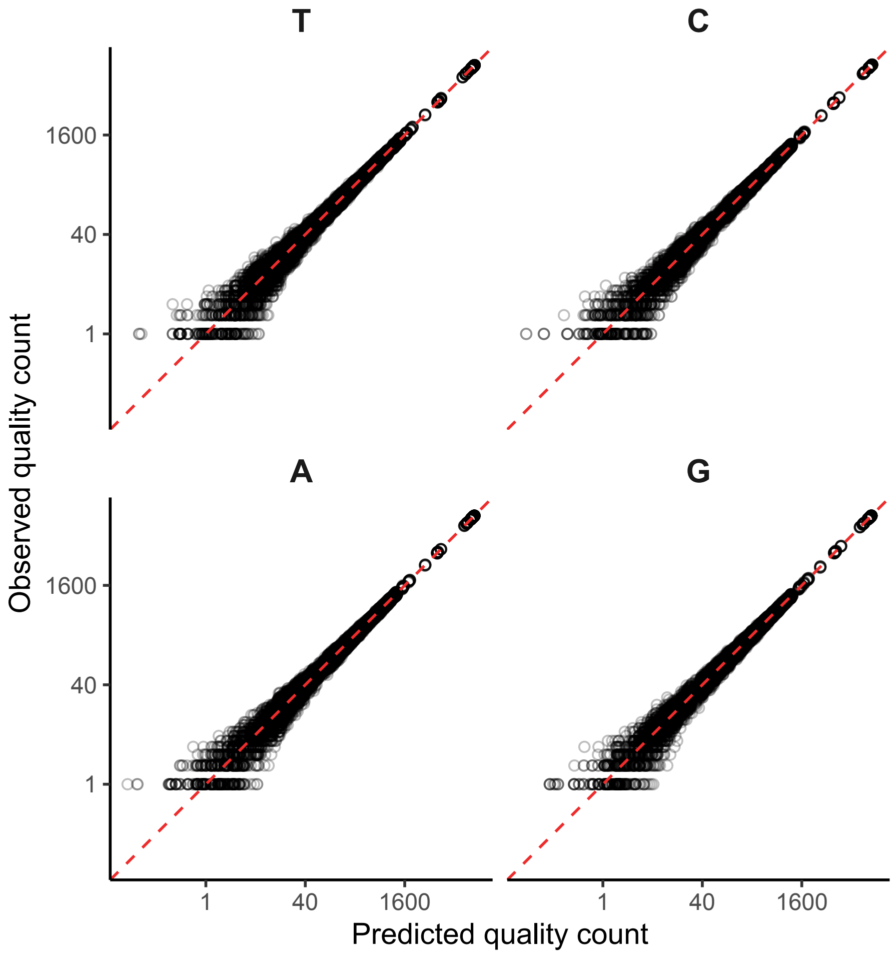

**Figure S8:** Observed versus predicted quality scores for each nucleotide. The dashed lines are the 1-to-1 lines. Each point represents the number of times a quality score was present at a given position and nucleotide for each of the 10 simulations.

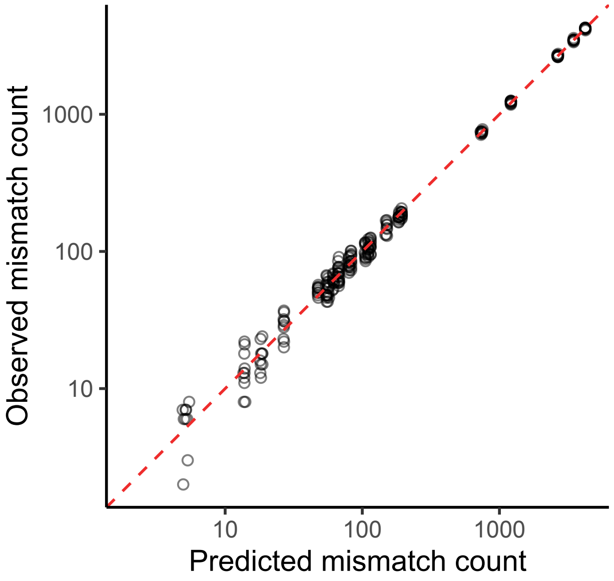

**Figure S9:** Observed versus predicted mismatches. The dashed line is the 1-to-1 line. Each point represents the number of mismatches for a given quality, for each of the 10 simulations.

#### 2.2 Testing indel rates

Indels are simulated the same as in ART, where there is a standard rate at which indels occur that does not depend on the position on the read. I am testing the accuracy of the output from `illumina` by starting with a reference genome with many small (smaller than the read length), same-sized sequences:

```
## < Set of 100,000 chromosomes >
## # Total size: 1,000,000 bp
##   name                          chromosome                           length
## chrom0     TATATATATA                                                    10
## chrom1     TATATATATA                                                    10
## chrom2     TATATATATA                                                    10
## chrom3     TATATATATA                                                    10
## chrom4     TATATATATA                                                    10
## ...        ...                                                          ...
## chrom99996 TATATATATA                                                    10
```

```
## chrom99997 TATATATATA                                                    10
## chrom99998 TATATATATA                                                    10
## chrom99999 TATATATATA                                                    10
```

Then I simulated 100-bp reads using a fake profile with no mismatches. Indels were found when the read length differed from the sequence length, and because I tested insertions and deletions separately, there was no chance of multiple indels offsetting one another. The predicted number of reads with at least one indel was simply the product of the indel rate, read length, and number of reads.

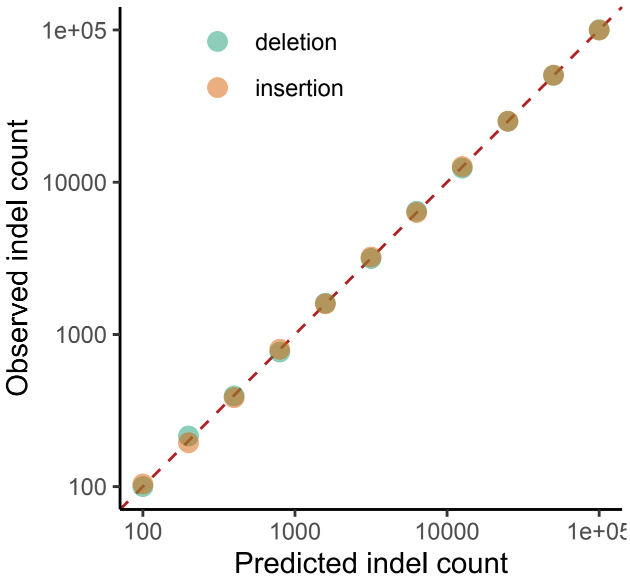

**Figure S10:** Observed versus predicted number of reads with indels. The dashed line is the 1-to-1 line. Note that all points overlap.

### 3 PacBio

This tests that the PacBio sequencer produces quality profiles and read lengths similar to expectations.

#### 3.1 Read lengths - default parameter values

For this section, I simulated 100,000 PacBio reads under default parameters. I then compared the read lengths to the expected read-length distribution:

$$\log(L - \delta) \sim \mathrm{N}(\mu, \sigma)$$

for read lengths $L \geq 50$, where $\delta = 10075.44$, $\mu = 9.79$, and $\sigma = 0.20$.

I used the following reference genome:

```
## < Set of 8 chromosomes >
## # Total size: 800,000 bp
##   name                          chromosome                          length
## chrom0     TAAACTATACCCGGGGGTTTTTAGG...AGGCGAGTAACCACAAAGCATATCAT    100000
## chrom1     CCTGATGCCGTGACAGACCCTTCAC...TTTACGTAACCACTACAGCCATAGGA    100000
## chrom2     CCCCTCAATTCAGGTTGACTGGCTT...AGCCACTTCACAGCGGTTCTAGAGTT    100000
## chrom3     AATGCGCGTTCGACCGTCGAGAAAA...AACCCGTGGGTGATCTGCTTTAGTGG    100000
## chrom4     TGCCTGCAAAGCGAGAGGAGGGAAT...GTTCCTAGGATCATAGAAAACGGACG    100000
## chrom5     TGTAAGACCAACATCGTGGAGGGCC...AGACATTTGCTCGTTCATGTCCGTGA    100000
## chrom6     CCTCGTATGGAGACTCTTTAGATGG...AGATGCATTATGTGAGTGCGTGAGCG    100000
## chrom7     CTATTCGACTATGTCCCCTCGCACG...GGGCGGCCAAAAAGCGTACTTACTAC    100000
```

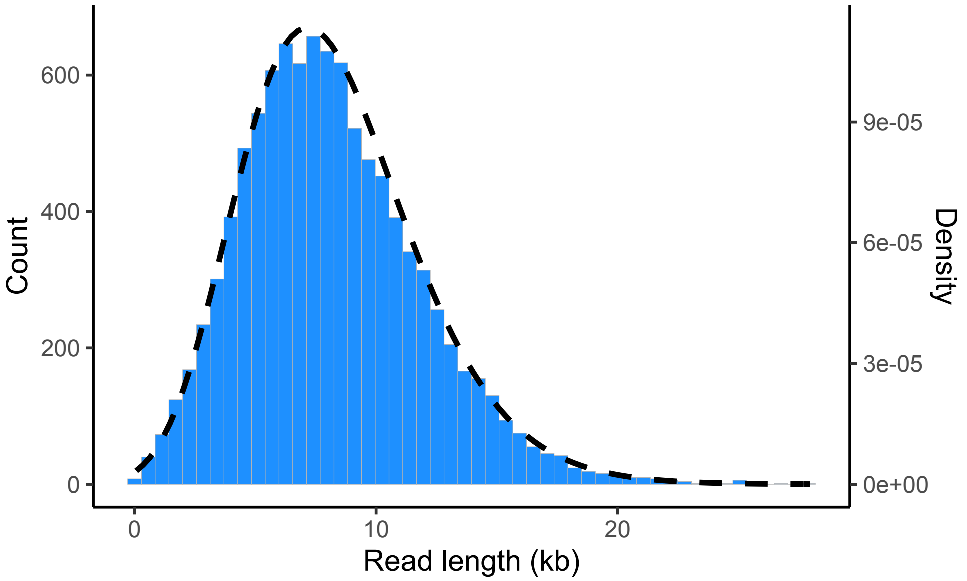

**Figure S11:** Read length distribution observed from 100,000 generated reads (histogram; left y-axis), alongside the frequency distribution as predicted for default parameters (dashed line; right y-axis).

#### 3.2 Read lengths - custom distribution

In this case, we generated 100,000 PacBio reads with a custom read-length distribution. The possible read lengths for this distribution were in the sequence from 100 to 10,000 by increments of 100. Each read length had a sampling weight drawn from a uniform distribution. So for a vector of weights ($W$) corresponding to each length, the number of predicted reads of the $i^{\text{th}}$ length is simply

$$N \frac{W_i}{\sum W},$$

where $N$ is the number of reads.

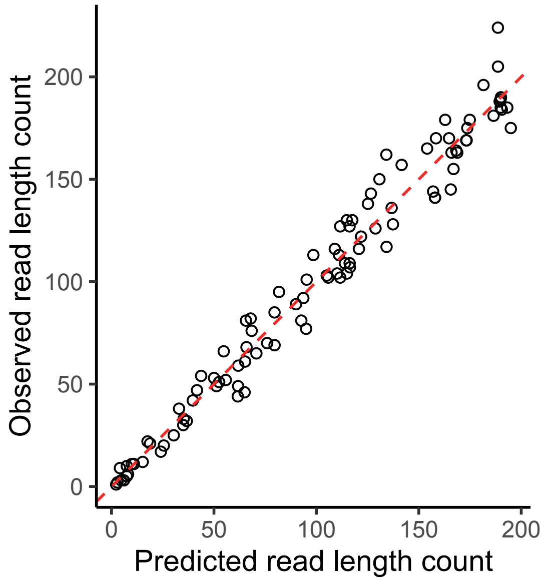

**Figure S12:** Observed versus predicted number of reads for PacBio reads generated from a custom distribution. The dashed line is the 1-to-1 line.

#### 3.3 Read qualities

Here I compared average read quality and read length between `jackalope` and `SimLoRD` by comparing the previous simulations performed in `jackalope` with output from the following `SimLoRD` command:

```
simlord --read-reference ref.fa -n 10000 --no-sam simlord_out
```

where `ref.fa` is a FASTA file written from the reference used in the `jackalope` simulations.

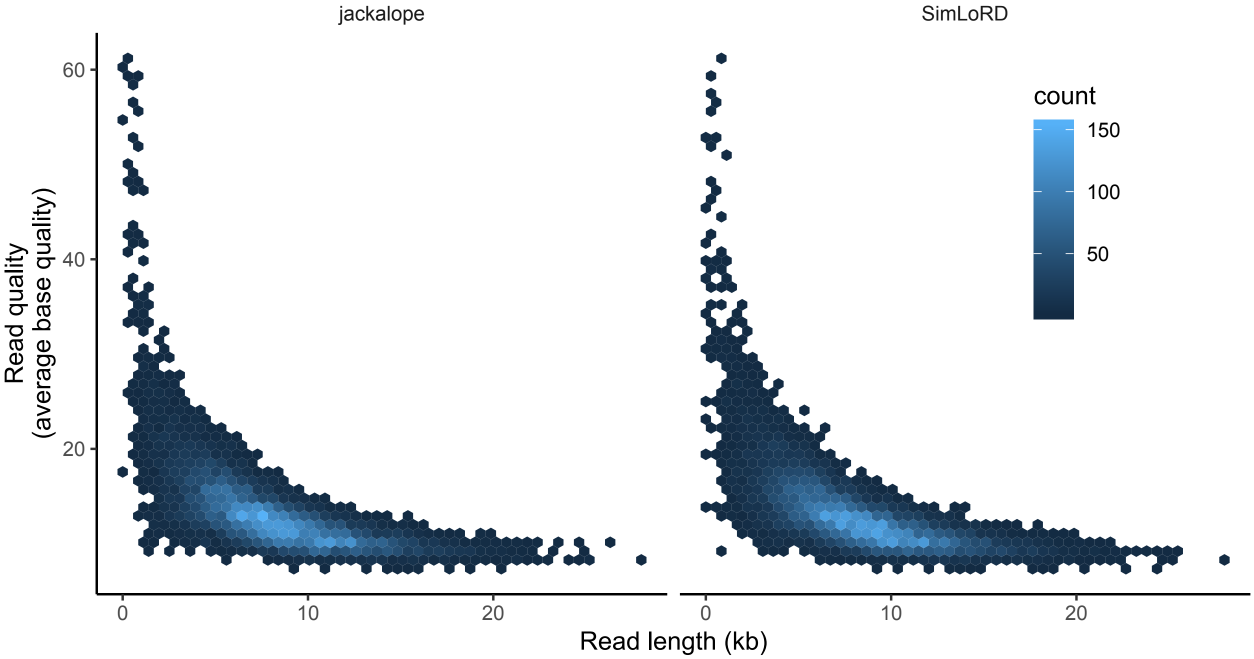

**Figure S13:** Read length and average quality per read for 10,000 PacBio reads simulated using `jackalope` and `SimLoRD`. Color of hex bins indicates the number of occurrences of that combination of read length and average quality.
